# Hierarchical Representation Learning for Drug Mechanism-of-Action Prediction from Gene Expression Data

**DOI:** 10.64898/2026.02.03.703502

**Authors:** Nikoletta Katsaouni, Marcel H. Schulz

## Abstract

Deciphering drug mechanisms of action (MoAs) from transcriptional responses is key for discovery and repurposing. While recent machine learning approaches improve prediction accuracy beyond traditional similarity metrics, they often lack biological structure and interpretability in the learned space. We introduce a hierarchical representation learning framework that explicitly enforces mechanistically coherent organization using dual ArcFace objectives, yielding an interpretable latent space that captures both MoA-level separation and compound-level substructure. Gene importance and pathway enrichment analyses confirm that the learned representations recover established signaling programs. Trained on LINCS L1000 data, the model also improves F1 performance over state-of-the-art baselines and generalizes to unseen compounds and cell types. Additionally, the latent space generalizes to CRISPR knockdowns without the need for retraining, indicating it captures pathway-level perturbations independently of modality.

## Introduction

Understanding the mechanism of action (MoA) of small molecules is central to drug discovery (Trapotsi et al., 2022) as it elucidates therapeutic mechanisms, enables repurposing, and guides the development of safer, more effective treatments (Pushpakom et al., 2019; Santos et al., 2017; Kibble et al., 2015).

Numerous computational approaches have been developed to predict MoA using different types of input data (Del Real and Rubio, 2023; Jang et al., 2021). Some of them rely solely on chemical structures such as molecular fingerprints or SMILES representations (Wu et al., 2018; Yang et al., 2019) to relate compounds through shared features or predicted molecular targets (Hosseini-Gerami et al., 2023; Liu et al., 2022). While structure-based models capture biochemical relationships between compounds, they provide limited insight into the functional consequences of drug treatment on cellular systems. Beyond these, knowledge-driven approaches — such as those leveraging molecular interaction networks, target annotations, or literature-derived associations — can infer pharmacological similarity but still depend heavily on prior knowledge of targets and pathways (Cousins et al., 2024).

In contrast, transcriptomic-based approaches use gene-expression profiles to characterize the downstream effects of perturbation, offering a direct view of how compounds reprogram cellular pathways and regulatory networks (Jiang et al., 2024). This perspective enables the identification of mechanistic similarities among compounds that may be chemically dissimilar and reveals context-dependent effects often invisible to structure-based predictions. The development of large-scale resources such as the Library of Integrated Network-based Cellular Signatures (LINCS) project (Subramanian et al., 2017), together with other efforts like the ChemPert (Zheng et al., 2023) have further accelerated this field by providing millions of standardized transcriptional signatures across diverse perturbations and cell types, paving the way for systematic, data-driven discovery of drug mechanisms.

From the approaches described above, most transcriptomic-based methods rely on one of two main strategies to quantify the similarity between gene expression signatures. Enrichment-based approaches, such as Gene Set Enrichment Analysis (GSEA) (Subramanian et al., 2005), compare the distribution of differentially expressed genes from a query compound to predefined gene sets, identifying coordinated pathway — a strategy recently extended by tools like the drug mechanism enrichment analysis (DMEA) (Garana et al., 2023) for mechanism-centric drug grouping. Other methods aggregate large collections of perturbation signatures (chemical and genetic) in unified repositories such as the iLINCS (Pilarczyk et al., 2022), which enables global signature similarity or connectivity analysis for drug repurposing and MoA inference. While effective for pathway-level interpretation, these methods depend on prior gene annotations and thresholds for up- and down-regulation, making results sensitive to reference quality and dataset variability. Distance-based methods instead compute similarity between transcriptional profiles using metrics such as Euclidean distance, cosine similarity, or rank-based correlation (Garana et al., 2023). By treating genes as independent features, these methods overlook gene–gene dependencies and nonlinear interactions, limiting biological interpretability and robustness.

Machine-learning approaches on the other hand have been used to learn latent representations that model nonlinear transcriptional relationships. Some methods focus on general perturbation similarity, embedding compounds based on distances in expression space without explicit MoA supervision (Yang et al., 2022; Cha et al., 2014). More recent contrastive frameworks, such as MOASL (Jiang et al., 2024), incorporate MoA labels to improve discrimination by pulling together signatures from the same mechanism. However, these embeddings primarily enforce separation at the MoA level and do not preserve meaningful substructure among individual compounds. Consequently, they offer limited support for downstream pharmacological analyses such as dose-dependent trajectory mapping, off-target exploration, or compound-level heterogeneity within a shared MoA.

In parallel, GPAR (Gao et al., 2021) formulates MoA prediction as multiple one-vs-rest classifiers, which prevents modeling global relationships among mechanisms and exacerbates class-imbalance issues common in LINCS data. Moreover, both representation-learning and binary-classification approaches remain sensitive to weak or noisy transcriptional responses.

To address these challenges, we developed a hierarchical representation learning framework that predicts MoA while organizing expression signatures into a biologically coherent latent space. Compounds with shared mechanisms cluster together, whereas individual drugs form meaningful substructure. The model disentangles pharmacological signals from cell-type variation and produces latent trajectories that reflect treatment strength.

Beyond strong predictive performance, the learned space is mechanistically interpretable: gene-level attribution and pathway enrichment highlight biologically relevant drivers of each MoA. The framework also generalizes to CRISPR perturbations without retraining, demonstrating robustness across modalities. Overall, our approach combines accurate classification with a structured embedding that supports downstream applications such as drug repurposing and mechanism discovery.

## Methods

### Dataset and Preprocessing

Transcriptional perturbation profiles were obtained from the LINCS L1000 project (Subramanian et al., 2017). The assay measures 978 landmark genes and infers additional genes using models trained on large-scale transcriptomic data; we used both measured and inferred genes to capture a comprehensive transcriptional response (Jiang et al., 2024). We used Level-5 consensus signatures, where each signature corresponds to a compound–cell line–dose–time condition and is defined as the median z-score differential expression across replicates. Data were downloaded from CLUE (https://clue.io/). Each signature is associated with a Transcriptional Activity Score (TAS) summarizing the number of differentially expressed genes and replicate consistency (Subramanian et al., 2017). Following Huang et al. (2025), we retained signatures with TAS ≥ 0.2 and discarded compounds with fewer than 10 signatures. MoA labels were taken from Huang et al. (2018), consistent with Jiang et al. (2024). After filtering, 28,769 signatures from 1,214 compounds were available for model development.

### Dataset Splitting

To avoid information leakage—an essential concern in drug repurposing as discussed in (Bernett et al., 2024)—and to obtain realistic estimates of model generalization, we created splits in which either compounds or cell types were entirely unseen during training.

- **Compound-based splits**. For each MoA, 20% of compounds were randomly assigned to the test set and the remaining 80% to training/validation, ensuring that test signatures come from entirely unseen drugs within known MoAs.
- **Cell-type–based splits**. To evaluate transfer to new cellular contexts, all signatures from a given cell line were held out for testing while the model was trained on the remaining cell lines. The held-out lines were NCIH508 (colorectal adenocarcinoma; 26 signatures), A549 (lung carcinoma; 2,209 signatures), MDAMB231 (triple-negative breast cancer; 646 signatures), and LNCaP (prostate adenocarcinoma; 94 signatures), representing diverse tissue origins.

These complementary regimes assess generalization across both chemical and biological space while avoiding compound-level overlap between train and test sets.

### Model Architecture and Training

A hierachical representation learning model was implemented in PyTorch to jointly capture the mechanism-level and compound-level structure from the gene expression signatures. The network consists of a fully connected encoder followed by two ArcFace-based classification heads corresponding to the MoA and compound respectively (Fig. 1).

**Fig. 1.**
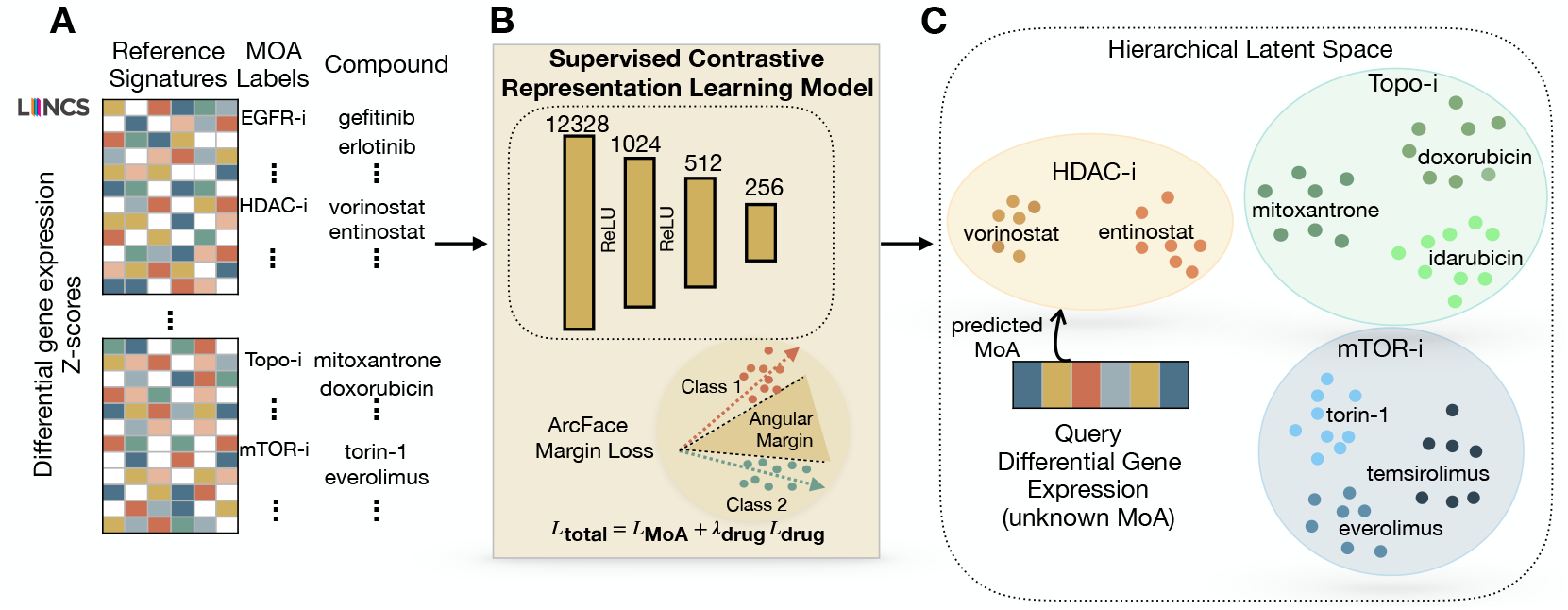
Overview of the hierarchical framework for predicting mechanisms of action (MoAs) from differential gene-expression signatures.(A) Reference LINCS Level-5 signatures, defined as drug-induced z-scores of the differential gene-expression profiles, are paired with known MoA annotations and used to train the model.(B) The model learns latent representations optimized with an ArcFace angular-margin loss, which enforces separation between MoA classes by maximizing inter-class angular distance while maintaining intra-class compactness.(C) In the resulting hierarchical latent space, compounds sharing the same MoA form distinct clusters with drug-level substructure. Query signatures with unknown MoA are projected into this space, and their mechanisms are inferred by similarity to reference clusters.

The fully connected network comprises three sequential linear layers with ReLU activations and a 256-dimensional latent embedding. The architecture was chosen after experimentation to balance between representational capacity and stability across the different folds.

#### ArcFace Loss Function

To impose a hierarchical organization in the latent space, we adopt the ArcFace loss (Deng et al., 2019), which optimizes class separation in angular space. For a normalized embedding **x**_*i*_ of sample *i* with ground-truth label *y*_*i*_, and a normalized class weight vector 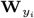,

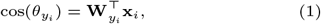

where 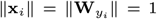. The index *i* iterates over samples in the mini-batch, and 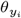 denotes the angle between the embedding and its class prototype.

ArcFace introduces an additive angular margin to enforce class separation. We extend this to a *class-specific* margin 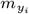 based on biological coherence: MoAs with high intra-class correlation, quantified as the mean pairwise Pearson correlation between all signatures belonging to that MoA, and sufficient support across all the folds are assigned a larger margin (0.70), while more heterogeneous or weakly supported MoAs receive a smaller margin (0.30). This strategy applies a larger margin only to MoAs that show strong and consistent transcriptional similarity, promoting clearer separation between biologically well-defined mechanisms. For more heterogeneous MoAs, a smaller margin prevents forcing signatures apart when their biological variability is naturally high.

The modified target logit becomes:

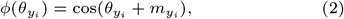

and the loss is computed using a scale factor *s* = 30:

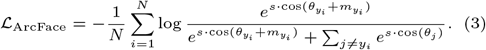

Our hierarchical formulation employs two cooperative ArcFace heads:

1. A MoA-level ArcFace head with class-specific angular margins 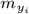.
2. A compound-level ArcFace head (margin = 0.1) that groups replicate profiles of the same drug.

The total training objective is:

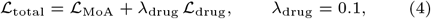

which jointly encourages MoA-level separation and drug-specific substructure, yielding a biologically organized latent space with interpretable pharmacological geometry.

#### Training Procedure

We optimized all parameters with Adam (learning rate 10^−5^, batch size 512) for up to 500 epochs, using early stopping on the validation loss. Each fold was trained independently on a single GPU. The resulting embeddings formed compact MoA clusters with drug-level substructure, capturing often dosage effects.

### Evaluation

The model performance evaluation was done on the hold-out test sets, as defined for each fold. We quantified intra-class similarity for each MoA using the mean pairwise Pearson correlation between all signatures assigned to that MoA. MoAs with higher intra-class similarity (*>* 0.1) under this metric were retained for evaluation. Details on the exact correlation values and the number of supports per MoA and per fold can be found in Supplementary File 1. The trained model was used to compute the latent embeddings for both training and test signatures through forward propagation. The predictive performance was inferred for the test compounds using a nearest-neighbor classifier operation in the learned latent space. Specifically the test compound’s representation was compared with those of training compounds to identify the most similar MoA. Additionally to the overall performance per fold, per-MoA metrics were computed for the estimation of mechanism-based performance variability.

#### Baseline Model Implementations and Evaluation

To contextualize the performance of our hierarchical ArcFace model, we compared it against three predictive baselines and a random classifier, all trained and evaluated on the same compound- and cell-type–held-out splits and using the identical LINCS Level-5 signatures.

##### MOASL (Jiang et al., 2024)

We reimplemented MOASL, a supervised contrastive learning model designed to learn similarity embeddings of perturbation signatures for MoA prediction. The model consists of a fully connected encoder with three hidden layers (1024 → 512 → 256 neurons), using Tanh activations. During training, signatures are grouped into predefined triplets—anchor, positive (same MoA), and negative (different MoA)—and optimized with a triplet cosine-similarity loss with a margin constraint, encouraging signatures with the same MoA to cluster while pushing apart different MoAs. After training, embeddings of reference signatures serve as a lookup space, and MoA labels are assigned via cosine-similarity *k*-nearest neighbors, following the original evaluation protocol. This allows predictions to be made solely based on the learned latent structure.

##### GPAR binary classifiers (Gao et al., 2021)

The original GPAR framework is formulated as a binary classifier for individual MoAs. To obtain a multi-class classifier, we trained a separate GPAR-style network for each MoA in a one-vs-rest fashion, keeping the architecture consistent with the published model (fully connected layers input → 978 → 512 → 256 → 1 with ReLU activations and dropout 10%). Each binary model was optimized with Adam learning rate (10^−4^) using a BCE-with-logits loss and an additional *ℓ*_1_ penalty on the weights for up to 2000 optimization steps per MoA. At test time, all binary networks were applied to each signature to obtain per-class sigmoid scores. The multi-class predictions were derived by taking the argmax over MoAs (with a score threshold of 0.5), and standard one-vs-rest metrics were computed for each mechanism.

##### Random Forest

As a strong baseline directly operating on gene-expression features, we trained a multi-class Random Forest classifier on the same input signatures. For each fold, we fitted a forest with 500 trees, class-balanced sample weights and default depth and splitting criteria on the training set. The model was evaluated on the corresponding held-out test set and the overall performance as derived from the class-probability outputs was reported.

##### Random baseline

Finally, we included a random baseline that assigns MoA labels to each test signature by sampling from the empirical MoA distribution observed in the training data. This classifier provides a lower bound on performance under our class imbalance and split strategy, with expected top-1 accuracy close to the inverse effective number of MoA classes.

### Cross-modality projection of CRISPR perturbation signatures

To assess modality-agnostic generalization, we evaluated whether the pretrained model could embed CRISPR-mediated gene perturbations consistently with drug-induced transcriptional responses. CRISPR knockdown signatures from the LINCS L1000 dataset were processed identically to small-molecule perturbation data. Briefly, we applied the frozen encoder to the normalized expression profiles to obtain latent representations, without any model retraining or fine-tuning.

MoA class probabilities were then generated using the pretrained MoA classifier, following the same inference procedure used for drug signatures. For illustrative examples, we selected target genes with well-defined associations to MoA categories present in the reference drug space and with sufficient experimental coverage. Specifically, MTOR was chosen as a canonical target of mTOR inhibitors, and CDK4 was selected as a representative cell-cycle kinase targeted by CDK inhibitors. Embedded CRISPR signatures were visualized alongside reference compound embeddings to evaluate cluster localization and mechanistic consistency.

### Interpretability

To explore the biological signals underlying the model predictions, an explainability analysis was performed on selected MoA classes. For each MoA a binary classifier was trained on top of the fixed pretrained network to distinguish compounds belonging to that MoA from all the others. This classifier consisted of a single linear layer trained using binary cross entropy loss while keeping the main pretrained network weights frozen.

Feature attribution was computed using Integrated Gradients (Sundararajan et al., 2017) to quantify each gene’s importance to the classifier output. The resulting gene contribution scores were averaged across samples, and the highest-ranking genes were interpreted as MoA-informative features according to our model.

To judge whether these genes correspond to biologically relevant processes, Gene Set Enrichment Analysis (GSEA) was applied to the ranked genes using the Reactome 2022 pathway collection as reference. The analysis was conducted with the GSEApy Python package (Fang et al., 2023), which implements the original Preranked GSEA algorithm (Subramanian et al., 2005). The identification of the enriched pathways (*FDR* < 0.05) permitted us to distinct between shared (core) and MoA-specific transcriptional programs. These analyses linked the model’s representations to mechanistic and pathway-level biological interpretation.

## Results

### Latent Space with Pharmacological Geometry

The model takes differential gene-expression signatures represented as z-scores, which quantify the transcriptional response to each compound. These signatures act as functional descriptors of drug action, and the model embeds them into a latent space designed to reflect underlying pharmacological relationships. The latent representation learned by the model revealed a biologically coherent organisation of compounds according to their MoA (Fig. 2A). The reduced predictive performance observed for certain MoA classes (for example channel blockers or serotonergic agents), as shown in Supplementary File 2, likely reflects intrinsic limitations of the training data rather than the model architecture. These compounds tend to induce weak or strongly cell-type–specific transcriptional responses that are not well captured by global z-score profiles, and they are represented by comparatively few, heterogeneous signatures. Consequently, the model has limited discriminative signal from which to learn these mechanisms. For this reason, we concentrated our quantitative analysis and benchmarking on MoAs that pass minimal support and intra-class correlation thresholds, where the available data are sufficiently rich to support mechanistically meaningful evaluation. Within this focused set, drugs with shared mechanisms formed compact and well-separated clusters in the latent space, indicating that the model successfully captures consistent transcriptional programs underlying pharmacological activity. Interestingly, compounds with related signaling effects, such as kinase inhibitors (PI3K-i, MEK/ERK-i, CDK-i) (Li et al., 2022; Chambard et al., 2007), occupied near regions of the space, suggesting that the embedding reflects functional proximity across pathways.

**Fig. 2.**
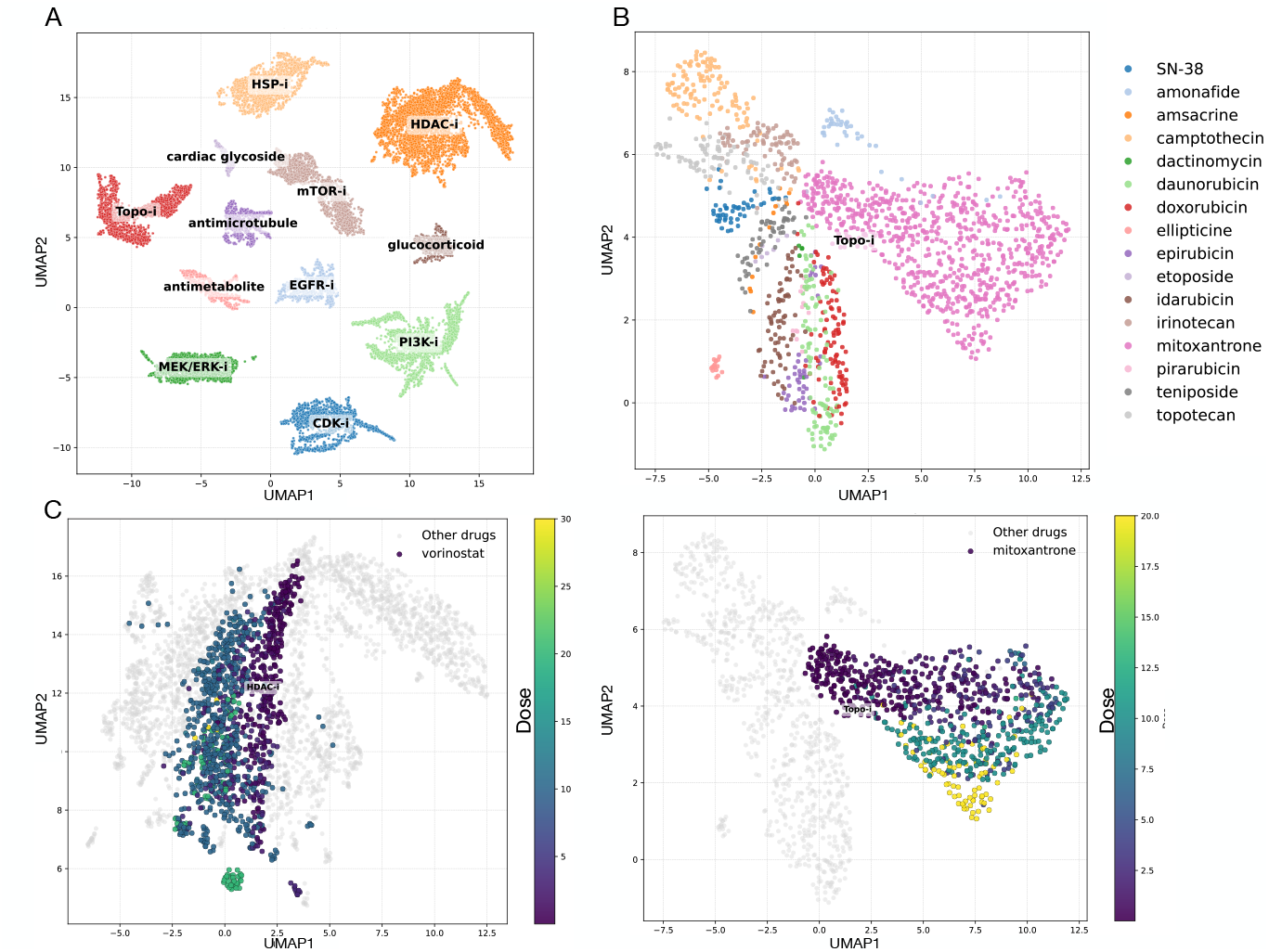
Hierarchical organization of the learned embedding space. (A) Compounds cluster according to their MoA in the learned latent space, where each point represents a transcriptional signature. (B) Within the Topoisomerase inhibitor MoA, individual drugs form distinct subclusters, reflecting subtle transcriptional differences. (C) Example compounds (mitoxantrone, vorinostat) with different MoAs form smooth dose-dependent trajectories, where the color bar indicates increasing compound concentration. This indicates that the latent space encodes pharmacodynamic variation.

Within individual MoA clusters, the latent structure preserved finer drug-level subclusters. As an example, in Fig. 2B the topoisomerase inhibitor subclusters correspond to specific compounds, such as doxorubicin, etoposide, and mitoxantrone. This organization indicates that the model not only groups modules by their shared primary target class but also captures secondary pharmacological differences and compound-specific response patterns.

Moving to the single compound level, replicate perturbations aligned along smooth trajectories across increasing doses (Fig. 2C), demostrating pharmacodynamic trends. This suggests that the model is able to embed both categorical and continuous aspects of drug response. Vorinostat and mitoxantrone were selected as representative examples because they display well-established, progressive transcriptional changes with dose (Munshi et al., 2006; Toh and Li, 2011), allowing clear visualization of how the embedding captures continuous variation in drug response.

### Model-derived gene importance reveals core stress responses and MOA-specific programs

To gain biological insight into the model’s representations, we analyzed the gene-level importance profiles derived from the explainability analysis (Fig. 3). For each MoA, the resulting Integrated Gradients scores provided a quantitative measure of how strongly each gene contributed to the model’s prediction, allowing us to identify the most informative genes for each drug mechanism.

**Fig. 3.**
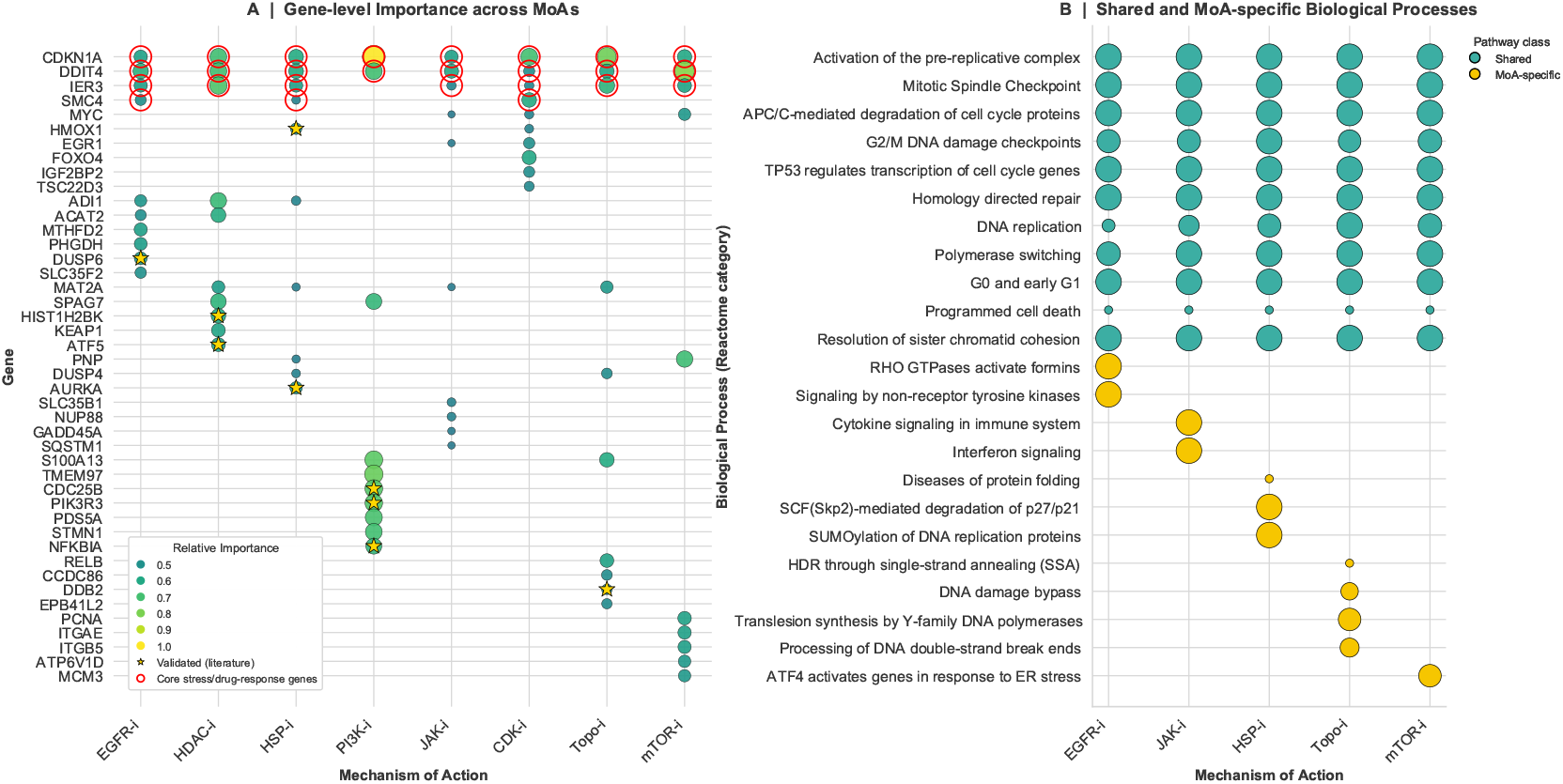
Gene- and pathway-level interpretation of model-derived features. (A) Gene importance across mechanisms of action (MoAs) based on Integrated Gradients. Bubble size and color represent per-MoA normalized gene importance (0–1 scaling, weighted by overall MoA magnitude). Red outlines indicate genes involved in broadly shared stress-response pathways, and stars denote literature-supported associations. (B) Reactome pathway enrichment of top-ranked genes. Pathways shared across drug-induced stress responses are shown in green, while MoA-specific signaling pathways are shown in yellow. Circle size is proportional to the mean − log_10_ nominal *p*-value across cross-validation folds, summarizing the strength and consistency of enrichment; larger circles indicate more significant pathway activation.

Because the MoA classifiers were trained independently, their raw importance scores were not directly comparable in magnitude. To address this, we applied a hybrid normalization strategy that preserved both within- and across-MoA interpretability. Specifically, gene importance values were first normalized between 0 and 1 within each MoA to retain the internal ranking structure, and then scaled by the overall magnitude of importance for that MoA. This weighting step ensured that differences in model calibration or dynamic range did not dominate the results, allowing the identification of biological patterns shared across mechanisms.

The resulting importance matrix revealed a coherent structure: a small set of genes showed high importance across nearly all MoAs, while others were uniquely enriched in specific drug classes. Among the shared genes, CDKN1A, DDIT4, IER3, and SMC4 consistently ranked among the top features across multiple MoAs. These genes are well-known regulators of cellular stress responses and growth control — CDKN1A (p21) mediates p53-dependent cell-cycle arrest (Engeland, 2022), DDIT4 acts as a stress-induced inhibitor of mTOR signaling (Jiao and Xiang, 2025), IER3 modulates MAPK and apoptotic signaling (Ye et al., 2018), and SMC4 contributes to chromatin remodeling and DNA damage repair (Pastic et al., 2024). The consistent prioritization of these stress-associated genes across distinct mechanisms indicates that the model captured a conserved core cellular stress and drug-response program, rather than spurious or MoA-specific noise.

In addition to these shared components, several MoA-specific signals emerged. For instance, PIK3R3 and STMN1 were among the top genes in the PI3K inhibitor (PI3K-i) class, consistent with their roles as downstream targets of PI3K signaling (Xun et al., 2021). Likewise, HMOX1 and AURKA were highlighted under heat-shock protein inhibition (HSP-i) (Barbagallo et al., 2015; Wang et al., 2022), while HIST1H2BK and ATF5 were specific to HDAC inhibitors (HDAC-i), reflecting their involvement in chromatin and transcriptional regulation (Alseksek et al., 2022). DUSP6 (a negative regulator of ERK signaling) and DDB2 (a DNA-damage-responsive protein) were identified under EGFR-i and Topoisomerase-i, respectively (Bagnyukova et al., 2013; Stoyanova et al., 2009). All the well documented associations are marked with a star in Fig. 3A), adding biological support to the model’s feature attributions.

Pathway-level enrichment (Fig. 3B) confirmed that the model captures biologically coherent structure in the latent space. Prominent shared pathways — including cell-cycle control, DNA replication/repair, and p53 signaling — reflect conserved stress responses induced by multiple cytotoxic MoAs, consistent with the central roles of CDKN1A and DDIT4 in growth arrest (Engeland, 2022; Jiao and Xiang, 2025). In contrast, PI3K/AKT signaling (PI3K-i), chromatin regulation (HDAC-i), ER-stress responses (HSP-i), and cytokine signaling (JAK-i) appeared as MoA-specific enrichments, in agreement with their known targeted mechanisms (Xun et al., 2021; Alseksek et al., 2022; Barbagallo et al., 2015; Wang et al., 2022; Bagnyukova et al., 2013; Li et al., 2022). Together, these results show that the model organizes signatures by mechanism while recovering established signaling biology. The full list of top enriched pathways for unseen-drug evaluation across all folds is provided in Supplementary File 3.

### Performance comparison with baseline and state-of-the-art models

We benchmarked our hierarchical representation learning model against MOASL, GPAR, a Random Forest baseline, and a random classifier. All methods were trained and evaluated on the exact same compound- and cell-type–held-out splits across all folds to ensure a fair and rigorous comparison of generalization performance.

Fig. 4 summarizes the evaluation. Across folds (Fig. 4A), our model achieved the highest mean F1 score (0.75), outperforming both recent deep learning approaches MOASL (0.73) and GPAR (0.73), while the Random Forest baseline lagged behind (0.64) and random assignment performed near zero. These results demonstrate that our hierarchical latent space enables more robust and accurate MoA prediction overall.

**Fig. 4.**
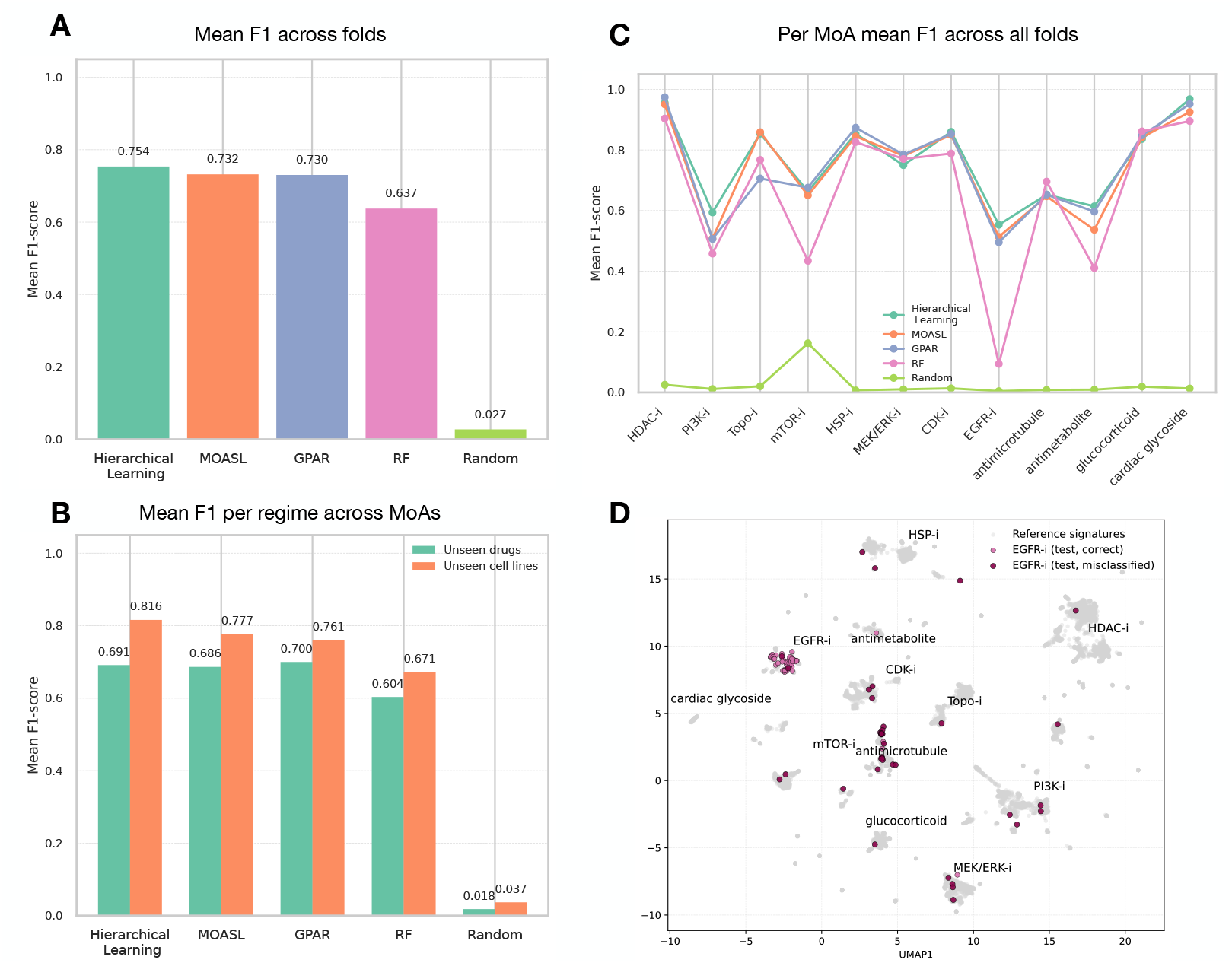
Evaluation of predictive performance under compound- and cell-type–held-out validation. (A) Mean F1 scores across all folds. (B) Performance comparison for unseen drug and unseen cell line regimes. (C) Mean per–mechanism of action (MoA) F1 scores. (D) UMAP visualization of the latent space for a representative fold, showing reference signatures (grey) and EGFR inhibitor test signatures (purple).

The hierarchical model also showed particularly strong performance when generalizing to unseen biological contexts (Fig. 4B). Although the three deep learning methods performed similarly when predicting MoAs for unseen drugs, our method achieved the best performance on unseen cell lines (mean F1 = 0.816), indicating that embedding compounds in a biologically structured space improves robustness under cell-line shifts.

Importantly, our method remains competitive or superior across nearly all MoAs retained after filtering for sufficient dataset support and meaningful intra-class transcriptional correlation. Detailed comparison across evaluation regimes (Table 1) shows that the proposed hierarchical learning model consistently achieves the highest mean F1 and recall, indicating more reliable identification of true MoA relationships. While GPAR attains strong ranking performance, our model maintains a more favorable balance between recall and precision, especially under the unseen cell-line regime. This balance is particularly important in discovery-oriented scenarios, where recovering true mechanistic effects is critical for advancing compounds rather than overlooking promising candidates. Together, these findings demonstrate that incorporating hierarchical structure in the latent space supports improved generalization across both chemical and biological variability, while retaining strong overall predictive accuracy.

**Table 1.** Performance comparison across models under compound- and cell-type–held-out evaluation. Results are reported as mean values across all folds. Best value per metric and regime is highlighted in **bold**.

| Model | Unseen Drugs |  |  | Unseen Cell Lines |  |  |
| --- | --- | --- | --- | --- | --- | --- |
|  | Precision | Recall | F1 | Precision | Recall | F1 |
| Hierarchical Learning | 0.66 | **0.75** | 0.69 | 0.77 | **0.84** | **0.82** |
| MOASL | 0.67 | 0.73 | 0.69 | 0.79 | 0.79 | 0.78 |
| GPAR | **0.79** | 0.66 | **0.70** | **0.87** | 0.73 | 0.76 |
| RF | 0.69 | 0.62 | 0.60 | 0.81 | 0.69 | 0.67 |
| RANDOM | 0.04 | 0.01 | 0.02 | 0.05 | 0.03 | 0.04 |

Per-MoA evaluation (Figure 4C) indicated that MoAs with well-differentiated transcriptional patterns, including HDAC, PI3K, and Topoisomerase inhibitors, were consistently classified by all models. To better understand why certain mechanisms remain challenging to classify, we focused on EGFR inhibitors (EGFR-i), which showed the lowest F1 score among the evaluated MoAs and for all the models. We visualized the EGFR-i test signatures in the learned latent space (Fig. 4D) to inspect how they organize relative to the reference clusters. Most EGFR-i signatures correctly mapped into the EGFR-i region, but a subset appeared closer to other MoAs. To further investigate the model’s context sensitivity, we examined predictions for two EGFR inhibitors with distinct pharmacological profiles: the FDA approved on-target drug afatinib and the more polypharmacologic investigational agent canertinib. As summarized in Table 2, afatinib is consistently classified as EGFR-i across multiple epithelial cell lines (Saber et al., 2016; Abourehab et al., 2021) and dose ranges, with errors restricted to extreme sub-effective or high-stress conditions. In contrast, canertinib exhibits systematic misclassification in EGFR-independent hematopoietic contexts (e.g. HBL1, TMD8, OCILY3) (Abourehab et al., 2021), where signatures resemble stress-associated mechanisms such as HSP-i, Topo-i or antimicrotubule agents. These results illustrate that misclassifications arise not always from random model failures, but from biologically grounded shifts in cellular response programs. The exact prediction results for all experimental conditions of afatinib and canertinib are provided in Supplementary File 4.

**Table 2.** Overview of model-derived MoA classifications for the EGFR inhibitors afatinib (FDA-approved) and canertinib (investigational) across representative cell lines and dosing conditions. This table presents representative predictions, indicating for each condition the cell line tested, the dose range (uM), whether the model correctly identified the EGFR-i MoA, the predicted MoA class, and a short description of the experimental context.

| Cell line(s) | Dose (uM) | Correct? | Predicted MoA | Interpretation |
| --- | --- | --- | --- | --- |
| **Afatinib (EGFR-i, FDA-approved)** |  |  |  |  |
| MCF10A, HPTEC, A549, SKBR3, LNCaP | 0.04–30 | Yes | EGFR-i | EGFR-dependent epithelial contexts |
| HA1E | 0.04 | No | CDK-i | Sub-effective dose |
| HPTEC | 100 | No | Topo-i | High-dose stress response |
| **Canertinib (EGFR-i, investigational)** |  |  |  |  |
| A549, A375, MCF10A, HPTEC | 0.04–10 | Mostly | EGFR-i | Effective only in EGFR-driven lines |
| HBL1, TMD8, OCILY3 | 0.3–10 | No | HSP-i / Anti-MT / Topo-i | EGFR-independent stress signatures |
| HT29 | 10 | No | HSP-i | Cytotoxic phenotype |

Together, these results indicate that structuring the latent space hierarchically—rather than learning only class-separating embeddings as in MOASL—provides measurable benefits: higher overall predictive accuracy, improved generalization to new biological contexts, and a mechanistically coherent space that supports downstream interpretability and exploratory drug analysis.

### Cross-modality validation on CRISPR perturbation profiles

To test whether the learned representation generalizes beyond chemical perturbations, we projected CRISPR-based gene knockdown signatures into the drug-derived latent space using the pretrained encoder and MoA classifier, without any retraining. We then visualized their positions relative to the reference drug clusters.

We examined two knockdown targets with direct mechanistic relevance to major MoA categories: MTOR for mTOR inhibition and CDK4 for CDK inhibition. In both cases, the CRISPR signatures were embedded jointly with the drug-induced profiles.

For MTOR knockdown, the CRISPR points localize directly within the mTOR-i cluster, overlapping with mTOR inhibitor compounds and receiving high prediction confidence for the mTOR-i MoA (Fig. 5, left). This demonstrates that the model interprets the transcriptional consequences of MTOR loss as comparable to pharmacological blockade.

**Fig. 5.**
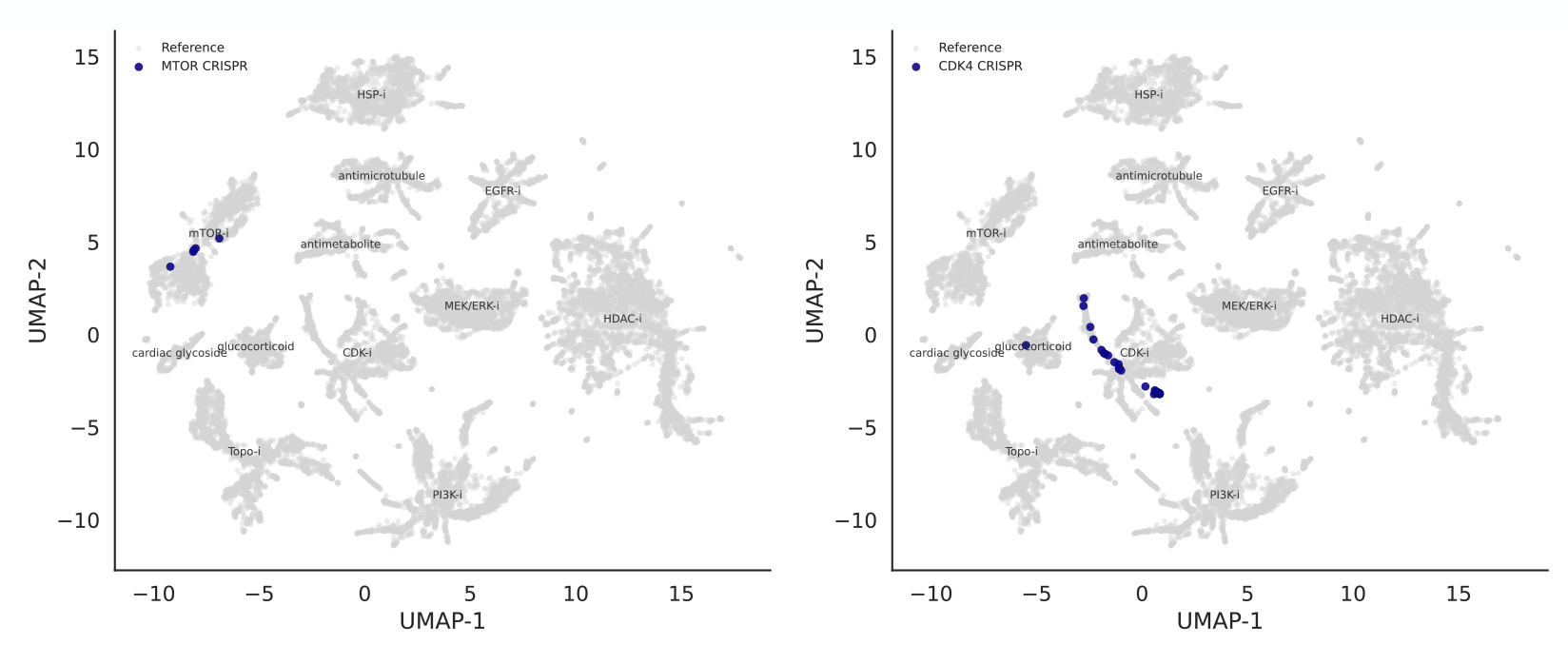
CRISPR perturbation signatures align with mechanistically corresponding drug clusters in the hierarchical latent space. Projection of gene knockdown transcriptional profiles (colored points) and drug reference signatures (gray points) into the learned latent space. Left: MTOR knockdown signatures cluster within the mTOR-i drug region and are assigned high probability for the mTOR-i MoA. Right: CDK4 knockdown signatures map into the CDK-i drug cluster, reflecting shared cell-cycle regulatory programs with pharmacological CDK inhibition.

Similarly, CDK4 knockdown signatures fall within the CDK-i neighborhood of the latent space, with consistently high CDK-i probabilities (Fig. 5, right). The trajectory formed by these profiles reflects a shared cell-cycle arrest program across genetic and chemical perturbations, while neighboring clusters representing distinct cytostatic mechanisms remain separate.

Together, these findings indicate that the hierarchical latent space organizes CRISPR perturbations in a manner coherent with drug MoAs, supporting the idea that the model captures pathway-level responses rather than perturbation-specific artifacts.

## Discussion and Conclusion

We introduced a hierarchical representation learning framework for predicting MoAs from transcriptomic perturbation data. Instead of treating MoA prediction solely as a multi-class classification task, our model explicitly structures the latent space using dual supervised heads: one enforcing MoA-level separation and one promoting compact drug-level organization. This yields clusters that reflect pharmacological similarity while preserving compound substructure and dose-dependent trajectories.

Across compound- and cell-held-out splits, our model matches or outperforms state-of-the-art methods such as MOASL and GPAR and a Random Forest baseline, with particularly strong gains on unseen cell types. The same representation supports downstream analyses including exploratory mapping of compound relationships, off-target hypotheses, and prioritization of repurposing candidates.

Interpretability analyses show that the model focuses on genes and pathways with known roles in stress response and MoA-specific signaling, linking the latent structure to established biology. Moreover, CRISPR knockdown signatures of MTOR and CDK4 map into the corresponding mTOR-i and CDK-i drug regions, suggesting that the representation captures pathway-level perturbations shared across modalities. To our knowledge, comparable MoA-prediction approaches have not previously demonstrated this capability using a single pretrained representation.

Our study also has certain limitations. Performance remains variable across MoAs, even after filtering out poorly supported classes. Some MoAs remain more difficult to classify because their transcriptional responses vary substantially across compounds and cellular contexts. Moreover, the model is trained on LINCS L1000 data and existing pharmacological annotations. Evaluating its robustness across additional assays and cell context will be an interesting next step.

Overall, this hierarchical framework strikes a practical balance between predictive accuracy and interpretability. It provides a biologically structured latent representation that generalizes across drugs, cell types, and perturbation modalities, offering a reusable backbone for applications in drug discovery and systems pharmacology—from systematic MoA inference to drug repurposing and integration of perturbation readouts such as CRISPR and proteomics.

## Data and Code Availability

The data used in this study is described in the manuscript and supplementary materials. The source code for data processing and analysis is publicly available at: [GitHub repository](https://github.com/SchulzLab/Hierarchical-Representation-Learning-for-Drug-MOA-Prediction)

---

GSEA pathway enrichment analysis for the 3 folds of unseen drugs for different MOAs

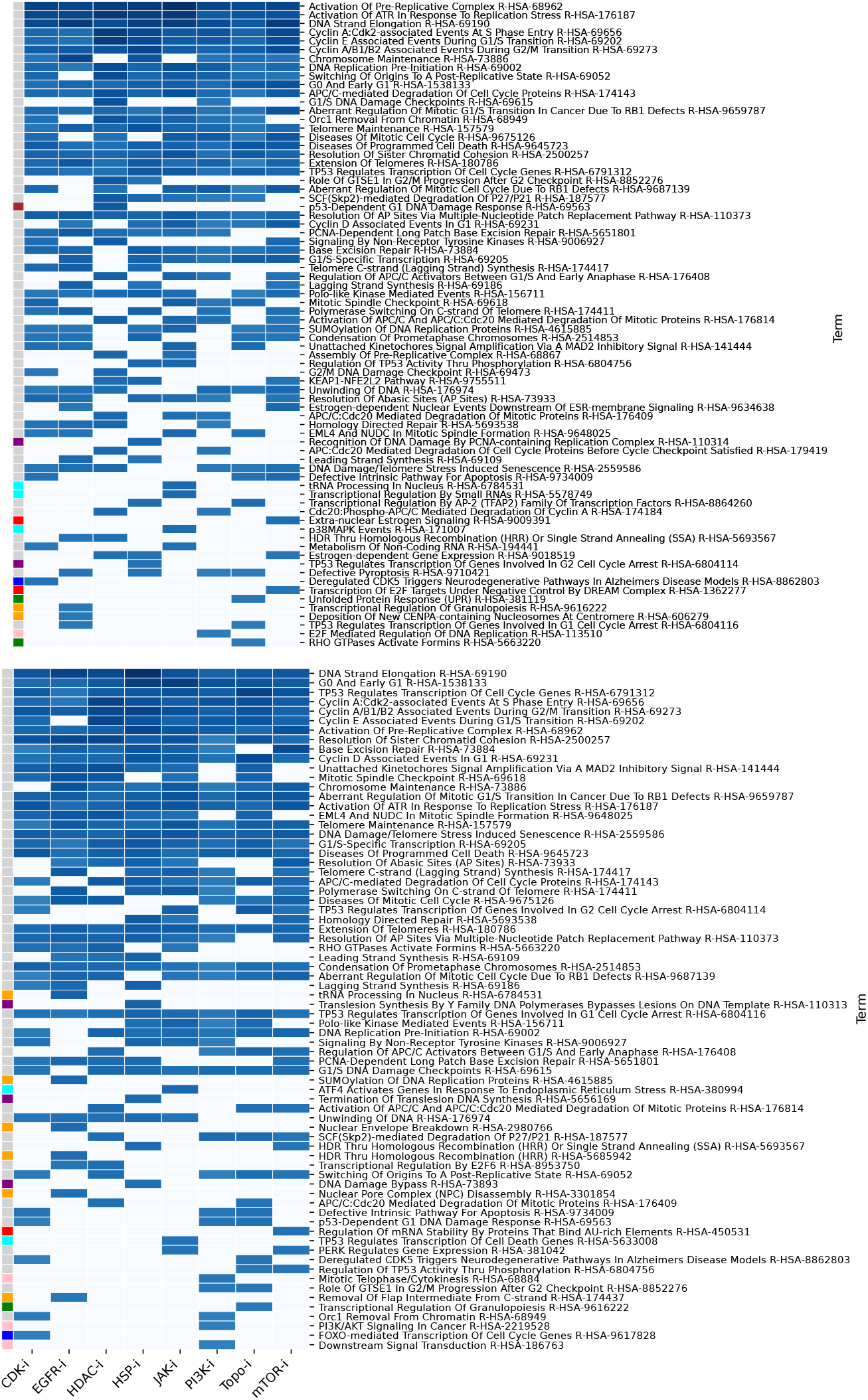
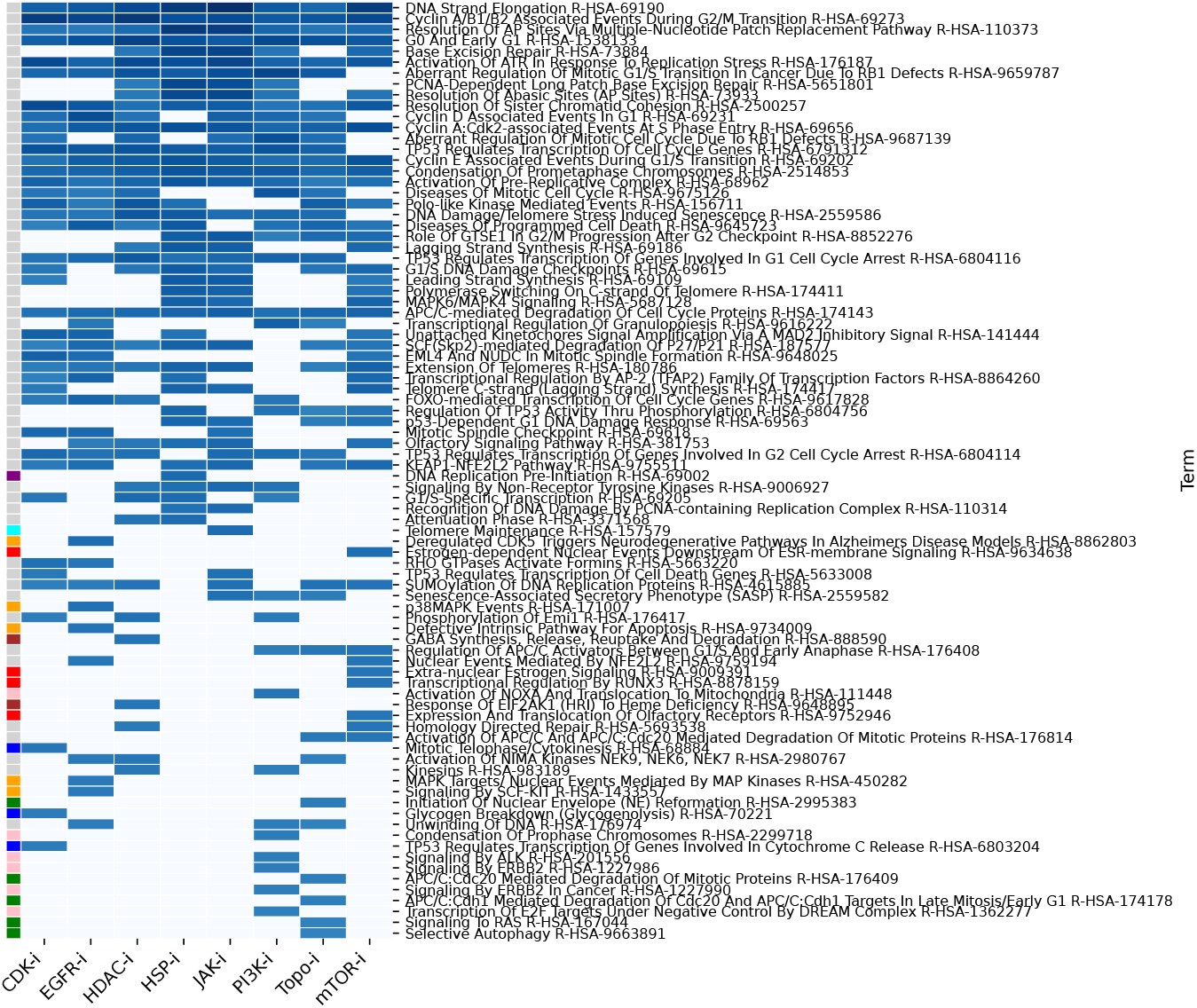

A UMAP visualization of all signatures for a single fold reveals that certain mechanisms of action are not well predicted for specific test samples.

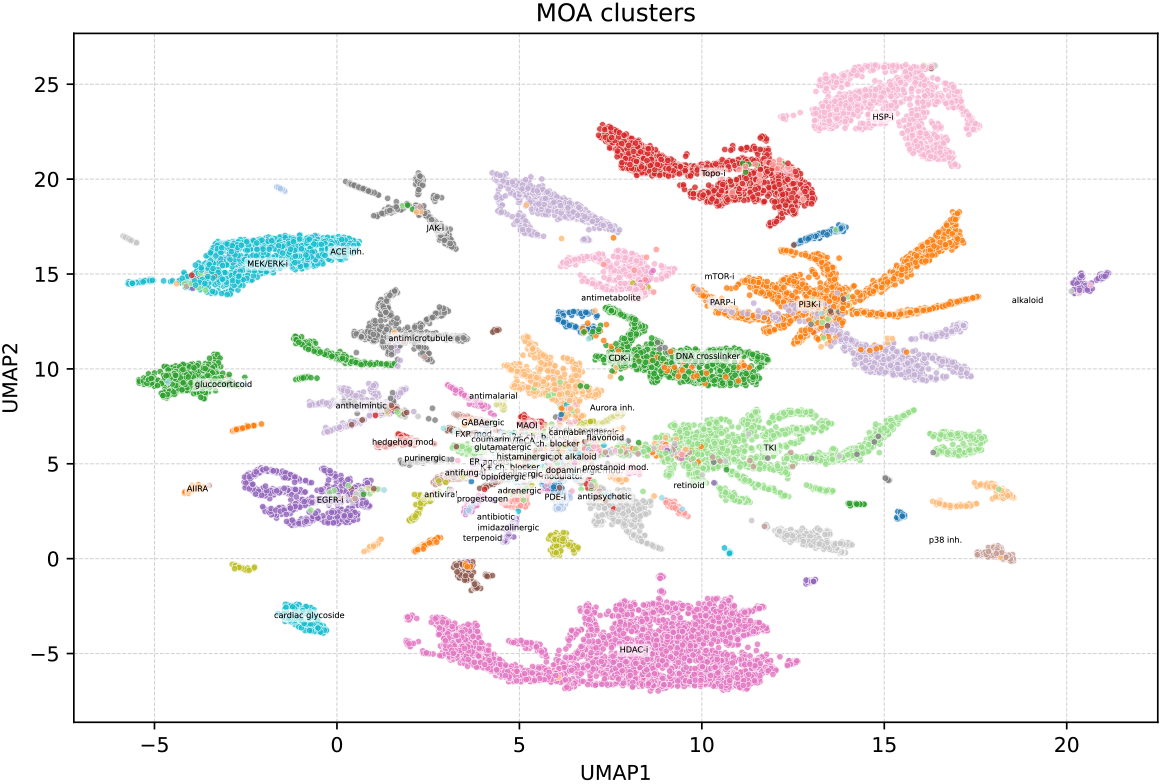

## References

M. A. Abourehab, A. M. Alqahtani, B. G. Youssif, and A. M. Gouda. Globally approved EGFR inhibitors: insights into their syntheses, target kinases, biological activities, receptor interactions, and metabolism. Molecules, 26(21):6677, 2021.

R. K. Alseksek, W. S. Ramadan, E. Saleh, and R. El-Awady. The role of hdacs in the response of cancer cells to cellular stress and the potential for therapeutic intervention. International Journal of Molecular Sciences, 23(15):8141, 2022.

T. Bagnyukova, D. Restifo, N. Beeharry, L. Gabitova, T. Li, G. Serebriiskii, E. Golemis, and I. Astsaturov. Dusp6 regulates drug sensitivity by modulating DNA damage response. British journal of cancer, 109(4):1063–1071, 2013.

Barbagallo, R. Parenti, A. Zappala, L. Vanella, D. Tibullo, F. Pepe, T. Onni, and G. L. Volti. Combined inhibition of Hsp90 and heme oxygenase-1 induces apoptosis and endoplasmic reticulum stress in melanoma. Acta Histochemica, 117(8):705–711, 2015.

J. Bernett, D. B. Blumenthal, D. G. Grimm, F. Haselbeck, R. Joeres, O. V. Kalinina, and M. List. Guiding questions to avoid data leakage in biological machine learning applications. Nature Methods, 21(8):1444–1453, 2024.

K. Cha, M.-S. Kim, K. Oh, H. Shin, and G.-S. Yi. Drug similarity search based on combined signatures in gene expression profiles. Healthcare Informatics Research, 20(1): 52–60, 2014.

J.-C. Chambard, R. Lefloch, J. Pouysségur, and P. Lenormand. Erk implication in cell cycle regulation. Biochimica et Biophysica Acta (BBA)-Molecular Cell Research, 1773(8): 1299–1310, 2007.

H. C. Cousins, G. Nayar, and R. B. Altman. Computational approaches to drug repurposing: methods, challenges, and opportunities. Annual Review of Biomedical Data Science, 7, 2024.

K. S. Del Real and A. Rubio. Discovering the mechanism of action of drugs with a sparse explainable network. EBioMedicine, 95, 2023.

J. Deng, J. Guo, N. Xue, and S. Zafeiriou. Arcface: Additive angular margin loss for deep face recognition. In Proceedings of the IEEE/CVF conference on computer vision and pattern recognition, pages 4690–4699, 2019.

K. Engeland. Cell cycle regulation: p53-p21-RB signaling. Cell Death & Differentiation, 29(5):946–960, 2022.

Z. Fang, X. Liu, and G. Peltz. GSEApy: a comprehensive package for performing gene set enrichment analysis in python. Bioinformatics, 39(1):btac757, 2023.

S. Gao, L. Han, D. Luo, G. Liu, Z. Xiao, G. Shan, Y. Zhang, and W. Zhou. Modeling drug mechanism of action with large scale gene-expression profiles using GPAR, an artificial intelligence platform. BMC bioinformatics, 22(1):17, 2021.

B. B. Garana, J. H. Joly, A. Delfarah, H. Hong, and N. A. Graham. Drug mechanism enrichment analysis improves prioritization of therapeutics for repurposing. BMC bioinformatics, 24(1):215, 2023.

L. Hosseini-Gerami, R. Hernansaiz Ballesteros, A. Liu, H. Broughton, D. A. Collier, and A. Bender. Maven: compound mechanism of action analysis and visualisation using transcriptomics and compound structure data in R/Shiny. BMC bioinformatics, 24(1):344, 2023.

C.-T. Huang, C.-H. Hsieh, Y.-J. Oyang, H.-C. Huang, and H.-F. Juan. A large-scale gene expression intensity-based similarity metric for drug repositioning. Iscience, 7:40–52, 2018.

K. Huang, P. N. Lalagkas, B. Sultan, and R. Melamed. Exploiting similarity in drug molecular effects for drug repurposing. Human Genomics, 19(1):110, 2025.

G. Jang, S. Park, S. Lee, S. Kim, S. Park, and J. Kang. Predicting mechanism of action of novel compounds using compound structure and transcriptomic signature coembedding. Bioinformatics, 37(1):376–382, 2021.

L. Jiang, S. Qu, Z. Yu, J. Wang, and X. Liu. MOASL: Predicting drug mechanism of actions through similarity learning with transcriptomic signature. Computers in Biology and Medicine, 169:107853, 2024.

Y. Jiao and Y. Xiang. A review of the participation of ddit4 in the tumor immune microenvironment through inhibiting PI3K-Akt/mTOR pathway. Frontiers in Oncology, 15: 1595463, 2025.

M. Kibble, N. Saarinen, J. Tang, K. Wennerberg, S. Mäkelä, and T. Aittokallio. Network pharmacology applications to map the unexplored target space and therapeutic potential of natural products. Natural product reports, 32(8):1249–1266, 2015.

Q. Li, Z. Li, T. Luo, and H. Shi. Targeting the PI3K/AKT/mTOR and RAF/MEK/ERK pathways for cancer therapy. Molecular biomedicine, 3(1):47, 2022.

C. Liu, A. M. Hogan, H. Sturm, M. W. Khan, M. M. Islam, A. Z. Rahman, R. Davis, S. T. Cardona, and P. Hu. Deep learning-driven prediction of drug mechanism of action from large-scale chemical-genetic interaction profiles. Journal of Cheminformatics, 14(1):12, 2022.

A. Munshi, T. Tanaka, M. L. Hobbs, S. L. Tucker, V. M. Richon, and R. E. Meyn. Vorinostat, a histone deacetylase inhibitor, enhances the response of human tumor cells to ionizing radiation through prolongation of γ-H2AX foci. Molecular cancer therapeutics, 5(8):1967–1974, 2006.

A. Pastic, M. L. Nosella, A. Kochhar, Z. H. Liu, J. D. Forman-Kay, and D. D’Amours. Chromosome compaction is triggered by an autonomous DNA-binding module within condensin. Cell reports, 43(7), 2024.

M. Pilarczyk, M. Fazel-Najafabadi, M. Kouril, B. Shamsaei, J. Vasiliauskas, W. Niu, N. Mahi, L. Zhang, N. A. Clark, Y. Ren, et al. Connecting omics signatures and revealing biological mechanisms with iLINCS. Nature communications, 13(1):4678, 2022.

S. Pushpakom, F. Iorio, P. A. Eyers, K. J. Escott, S. Hopper, A. Wells, A. Doig, T. Guilliams, J. Latimer, C. McNamee, et al. Drug repurposing: progress, challenges and recommendations. Nature reviews Drug discovery, 18(1): 41–58, 2019.

A. Saber, T. Hiltermann, K. Kok, H. Groen, et al. Resistance mechanisms after tyrosine kinase inhibitors afatinib and crizotinib in non-small cell lung cancer, a review of the literature. Critical reviews in oncology/hematology, 100: 107–116, 2016.

R. Santos, O. Ursu, A. Gaulton, A. P. Bento, R. S. Donadi, C. G. Bologa, A. Karlsson, B. Al-Lazikani, A. Hersey, T. I. Oprea, et al. A comprehensive map of molecular drug targets. Nature reviews Drug discovery, 16(1):19–34, 2017.

T. Stoyanova, N. Roy, D. Kopanja, P. Raychaudhuri, and S. Bagchi. Ddb2 (damaged-dna binding protein 2) in nucleotide excision repair and DNA damage response. Cell Cycle, 8(24):4067–4071, 2009.

Subramanian, P. Tamayo, V. K. Mootha, S. Mukherjee, L. Ebert, M. A. Gillette, A. Paulovich, S. L. Pomeroy, T. R. Golub, E. S. Lander, et al. Gene set enrichment analysis: a knowledge-based approach for interpreting genome-wide expression profiles. Proceedings of the National Academy of Sciences, 102(43):15545–15550, 2005.

A. Subramanian, R. Narayan, S. M. Corsello, D. D. Peck, T. E. Natoli, X. Lu, J. Gould, J. F. Davis, A. A. Tubelli, J. K. Asiedu, et al. A next generation connectivity map: L1000 platform and the first 1,000,000 profiles. Cell, 171(6):1437–1452, 2017.

M. Sundararajan, A. Taly, and Q. Yan. Axiomatic attribution for deep networks. In International conference on machine learning, pages 3319–3328. PMLR, 2017.

Y.-M. Toh and T.-K. Li. Mitoxantrone inhibits HIF-1α expression in a topoisomerase ii–independent pathway. Clinical cancer research, 17(15):5026–5037, 2011.

M.-A. Trapotsi, L. Hosseini-Gerami, and A. Bender. Computational analyses of mechanism of action (MoA): data, methods and integration. RSC chemical biology, 3(2): 170–200, 2022.

F. Wang, H. Zhang, H. Wang, T. Qiu, B. He, and Q. Yang. Combination of aurka inhibitor and HSP90 inhibitor to treat breast cancer with AURKA overexpression and TP53 mutations. Medical Oncology, 39(12):180, 2022.

Z. Wu, B. Ramsundar, E. N. Feinberg, J. Gomes, C. Geniesse, A. S. Pappu, K. Leswing, and V. Pande. MoleculeNet: a benchmark for molecular machine learning. Chemical science, 9(2):513–530, 2018.

G. Xun, W. Hu, and B. Li. PTEN loss promotes oncogenic function of STMN1 via PI3K/AKT pathway in lung cancer. Scientific Reports, 11(1):14318, 2021.

C. Yang, H. Zhang, M. Chen, S. Wang, R. Qian, L. Zhang, X. Huang, J. Wang, Z. Liu, W. Qin, et al. A survey of optimal strategy for signature-based drug repositioning and an application to liver cancer. Elife, 11:e71880, 2022.

K. Yang, K. Swanson, W. Jin, C. Coley, P. Eiden, H. Gao, A. Guzman-Perez, T. Hopper, B. Kelley, M. Mathea, et al. Analyzing learned molecular representations for property prediction. Journal of chemical information and modeling, 59(8):3370–3388, 2019.

J. Ye, Y. Zhang, Z. Cai, M. Jiang, B. Li, G. Chen, Y. Zeng, Y. Liang, S. Wu, Z. Wang, et al. Increased expression of immediate early response gene 3 protein promotes aggressive progression and predicts poor prognosis in human bladder cancer. BMC urology, 18(1):82, 2018.

M. Zheng, S. Okawa, M. Bravo, F. Chen, M.-L. Martínez-Chantar, and A. Del Sol. ChemPert: mapping between chemical perturbation and transcriptional response for non-cancer cells. Nucleic Acids Research, 51(D1):D877–D889, 2023.

